# A variant of the Morris water task for assessing learning and memory processes in mice

**DOI:** 10.1101/177105

**Authors:** Jogender Mehla, Jamshid Faraji, Takashi Saito, Takaomi C Saido, Majid H. Mohajerani, Robert J. McDonald

## Abstract

The Morris water task (MWT) is commonly used to assess rodent spatial learning and memory. Our goal was to develop a 3-phase variant of the hidden goal water task to assess old and new spatial memories acquired in the same context using various measures of spatial learning in C57BL/6 mice. In the first phase, mice were pre-trained to an initially hidden location. The second phase consisted of a massed training session to a new location in the same apparatus and context. The final phase consisted of a competition test between the original and new platform locations. *App*^*NL-G-F/NL-G-F*^ mice, a novel transgenic mouse model for Alzheimer’s disease (AD), were also used as an independent variable to validate this 3-phase variant of MWT. The results of the present study showed that C57 mice acquired and retained both the old and new location representations; however, *App*^*NL-G-F/NL-G-F*^ mice retained a recently acquired spatial memory but did not remember the old location acquired in the same apparatus and context. The results showed that C57 mice can show precise place learning and memory with the right amount of training and acquire and retain multiple spatial memory locations in the same environment whereas this ability was impaired in *App*^*NL-G-F/NL-G-F*^ mice. In the visible platform test, however, all groups of mice showed normal sensorimotor ability and motivation. These findings indicate that this new version of the MWT provides a robust way for assessment of old and new memories in mice. This paradigm could also be exploited to assess manipulations of neural circuits implicated in learning and memory processes as well as for research investigating human brain diseases.

## Introduction

Some consider a deep understanding of the organization of learning and memory functions in the mammal and the neurobiological processes underlying this fundamental brain function one of the great challenges facing the field of systems neuroscience^1-3^. One view suggests that this work will not only inform us about the neural systems and mechanisms underlying learning and memory processes but also how they contribute to normal manifestations of thought and behaviour^4^ and what happens when these systems are impaired in disease states like age-related cognitive decline^5^.

A central structure of one learning and memory system, the hippocampus, is synonymous with learning and memory function and decades of research supports the claim that this medial temporal lobe structure is part of a relational learning and memory system^6-8^ that supports episodic memory processes. The hippocampus and associated neural circuits that make up this complex learning and memory system have been shown to be essential when an animal has to learn about the relationships amongst cues to solve a particular task. Rats with hippocampal damage have been reported to be impaired at the acquisition and retention of various spatial tasks including the Morris water task (MWT)^9-11^. Evidence also indicates that rats with hippocampal damage are impaired at variants of context conditioning^12-14^ and other non-spatial relational tasks including various configurable association tasks^15^.

Of the many learning and memory tasks developed by behavioural neuroscientists, the MWT has proven to be the most useful and versatile assay of learning and memory function in rodents and a reliable and valid measure of hippocampal function^4^. The spatial version of the water task is one of the most widely used learning and memory tests for rodents. This popularity is well founded because the task has various design features that make it a powerful tool for assessing rodent learning and memory function. The features include the use of a pool that is completely homogeneous (no local cues). This ensures that the subjects cannot use intra-maze features to find the escape platform, thus requiring subjects to use distal cues in the testing room. If local features are present, then rats with hippocampal damage may achieve normal levels of performance^10^. A second important design feature of this task is the use of different start positions that are located at the four cardinal compass points (N, S, E, W). Because the starting positions are pseudo-randomly selected for each swim, it is difficult for subjects to navigate based on a simple turning response or guidance strategy such as approaching a prominent distal cue. Another key feature of the water task is that it utilizes a swimming behaviour that rats use in their natural ecosystem making the task a valid assessment of rat cognitive function. Another important design feature of the water task is the use of a non-invasive tracking system and computerized data collection which can reduce experimental bias. The task also allows for assessment of sensory, motor, and motivational functions to ensure that impairments found on the task are not due to disruption of these brain functions that support learning and memory functions but are not cognitive processes *per se*. These brain functions that support learning and memory are assessed by using visible (cued) platform training procedures and assessment of swimming abilities. Finally, the water task does not require food deprivation to motivate subjects and human analogues of the task have been developed for cross-species comparisons that integrate animal model data with human research^16^.

The MWT was originally designed to evaluate spatial learning and memory processes in rats but in recent years is also used very commonly for mice^17,18,11^. The goal of the present experiments was to develop a version of the three-phase MWT for use in mice. This was deemed an important missive as mice are an important model of the neural basis of learning and memory because of their dominant role in trying to understand the genetic basis of learning and memory as well as genetic contributions to various neurodegenerative disorders linked to age-related cognitive decline like Alzheimer’s disease (AD).

For the task design utilized in the present study, we were inspired by a 3 phase MWT paradigm developed for a set of behavioural pharmacology experiments in rats^19^. Briefly, phase 1 consisted of standard spatial training to a static hidden platform location until asymptotic levels of performance are achieved. For phase 2, the subjects are given mass training to a new location in the same apparatus and context over a single day. In the final phase, the subjects are given a competition test in which they are allowed to freely swim to either the new or old spatial location. The task was originally designed this way because of controversies surrounding the role of NMDA receptors in hippocampal learning and memory processes. Although still unresolved, one pattern of results that has emerged from this work is that NMDA receptor blockade does not seem to impair hippocampal spatial learning and memory if the rats are pre-trained on the MWT. The idea was that the initial reports showing the NMDA receptor blockade impaired acquisition of the MWT were due to sensory/motor/motivational impairments caused by the pharmacological manipulation, and not because plasticity processes supporting learning and memory had been compromised. Pre-training was thought to reduce these effects by allowing the subject to acquire the procedural aspects of the task before blockade of NMDA receptor plasticity. Our research group incorporated this pre-training procedure (phase 1) to ensure that any functional effects of our NMDA pharmacological manipulations were not due to non-mnemonic effects. It was during phase 2 that NMDA receptors were blocked. This allowed the assessment of the role of NMDA receptors on rapid acquisition of new information on the MWT in behaviourally-sophisticated subjects. Further, phase 3 gave the subjects the opportunity to exhibit a preference for the new or old spatial location, both of which provided escape from the cold water at some point in training. A clear demonstration of the influence of new versus old memories.

In summary, this task design is important because of the assessment of initial learning over many days which also provides important procedural experiences to the subject. A rapid mass training phase in which acquisition of new information in one day in a behaviourally sophisticated subject is obtained. Finally, an assessment of the influence of new versus old memories is revealed.

In the present study, we designed a modified version of the three-phase MWT which included a novel assessment of phase 2 rapid acquisition trials to determine how quickly the mice will learn the new location of the platform and when the spatial behaviour of the mice would shift from old memory to new memory control. Accordingly, we assessed the ability of adult male C57BL mice to learn in a three-phase variant of the MWT that included the acquisition of an original escape platform location over several days of training, rapid mass training for a new platform location, and a competition test to assess the relative influence of the new and old location representations. In the present study, we also assessed a newly developed single amyloid precursor protein knock-in mouse (*App*^*NL-G-F/NL-G-F*^) model for AD as an independent variable to validate this three-phase variant of the water task. The novel AD model in mice has been developed by Saito and colleagues to overcome the problem of over expression of Aß precursor protein (App) in transgenic mouse models for AD^20,21^. Moreover, *App*^*NL-G-F/NL-G-F*^ mice also showed the deficits in spatial memory and flexible learning, enhanced compulsive behavior, and reduced attention performance in previous study^22^.

## Materials and Methods

### Animals

Twenty-nine male mice were used in the present study. Eight C57BL and 13 C57BL mice (4-6 months old) were used in experiments 1 and 2, respectively. Also, 8 *App*^*NL-G-F/NL-G-F*^ mice (9 months old) were used in experiment 2 as an independent variable to validate this protocol. The male and female pairs of AD transgenic mice carrying Swedish (NL), Arctic (G), and Beyreuther/Iberian (F) mutations (*App*^*NL-G-F/NL-G-F*^) were provided by RIKEN Brain Science Institute, Japan. Then, a colony of these mice was maintained at the Canadian Center for Behavioural Neuroscience. Genotyping of all mice was done by PCR using tail snipping method. All animals were group-housed (4 per cage) and had food and water *ad libitum*. They were also maintained on a 12:12 light/dark cycle. Moreover, mice were individually handled each day (approximately 2-3 min) for one week before any experimental manipulation. All testing and training were performed during the light phase of the cycle at the same time of day. All experimental procedures were approved by the University of Lethbridge Animal Care Committee in compliance with the standards set out by the Canadian Council for Animal Care.

### Apparatus

The spatial performance was tested in MWT according to previous studies with some modifications^23,24^. The task consisted of a pool (154 cm diameter) filled to within 20 cm of the top of the wall with water (22±1°C) that was rendered opaque by non-toxic tempera white paint. The pool was located in a room rich with distinct distal cues, which remained unobstructed throughout the duration of the experiment. A circular escape platform (12 cm radius) was submerged 0.5-1 cm below the water surface and located at any one of four quadrants.

### Experiment 1

#### Phase one: original location training

Mice in this experiment received 4 training trials from each starting point in distributed manner for 8 days. During each trial, mice were placed into the pool, facing the wall, in one of the four start locations (North, South, East, and West). The trial was completed once the mouse found the platform or 60 sec had elapsed. If the mouse failed to find the platform on a given trial, the experimenter guided the mouse onto the platform. The latency (time to locate the hidden platform), distance travelled by mouse before reaching the platform, corridor percent path (error index)^25^ and Gallagher’s measurements (cumulative and average proximities)^26^ were analyzed to assess learning and memory processes in each mouse. A corridor (25 cm wide) was drawn between the start location and the platform for corridor percent path. The corridor percent path was also recorded during these trials to examine whether the mice swam in a straight path from the start location to the platform^27^. To understand the search strategy the animal uses during the trials, Gallagher et al. (1993) measured these proximities measurements which provide an idea about the distance between animal and platform. The swim speed was also analyzed to rule out the involvement of altered motor function as a confounding factor^26^. All measures of spatial navigation in the MWT were recorded and analyzed by an image-computerized tracking system (HVS Image 2020, UK).

#### Probe trial

A probe trial in which the escape platform was removed from the pool was conducted on day 9 as a measure of spatial memory. The platform in the probe trial was removed from the target quadrant (SE), and the trial was started with the mice being placed in the water tank from the north side. Then, the mice were allowed to swim freely for 60 sec. The data collected from the probe trial was analyzed by measuring the quadrants preference (percentage of time that the animals spent in each quadrant) and distance travelled in target quadrant. The data in probe trial was analyzed for 0-30 sec and 60 sec time bins.

#### Phase two: new location training (rapid mass training)

Phase 2 of training commenced on day 9 after the completion of the probe trial test. For the mass training, the hidden platform was moved to the northwest pool quadrant for new location training. For new location training, a total of 20 trials divided into 5 trial blocks were given on day 9 using all four starting points in distributed manner. The inter-trial time was approximately 15 to 20 min. In this phase, the data were analyzed by measuring the latency, distance travelled, corridor percent path, Gallagher’s measurement and assessment of preference for the old and new location quadrants.

#### Phase three: competition test

The competitive test was performed 24 hrs after phase 2 training was completed. The starting point for this probe trial was equidistant from old and new quadrants location of the platform. The data collected from the competition test was also analyzed by measuring the quadrants preferences and distance travelled in old and new location quadrants. The competition test would provide an idea about the influence of the new versus old location spatial memories on overt behavior and an indirect measure of the relative strength of the two hippocampal representations. In this phase, the data was analyzed for 0-30 sec and 60 sec bins. Moreover, we also analyzed the data in time bin manner like 0-10, 11-20 & 21-30 sec to investigate the quadrant preference of mice in the early, middle, and end components of the competition test. Previous work has shown that an early preference for the new platform position in rats that switches to a preference for the old location. This indicates that animals had both representations available to them during the competition test with the new spatial representation controlling behavior initially^19^.

### Experiment 2

#### Phase one: original location training

The protocol for acquisition phase learning was similar to experiment 1.

#### Probe trial

The protocol for probe trial was similar to experiment 1.

#### Phase two: new location training (rapid mass training)

Before starting this phase, we gave the mice 8 additional training trials to the old location quadrant to rule out the possible loss of memory during the probe trial. The protocol in this phase was similar to experiment 1 with the exception that training consisted of 32 trials divided into 8 trial blocks, which was equivalent to the normal acquisition phase were given to mice. The change in the number of training trials in this phase was based on the demonstration, in experiment 1, that the mice in experiment 1 did not show a preference for the new location during the competition test suggesting that the strength of the new location memory was not strong enough to gain control over behavior. We posited that more training during phase-2 would produce a memory for new location during the competition test.

#### Phase three: competition test

The protocol for the competition test was identical to experiment 1.

#### Visible platform test

Performance in the MWT may be affected by deficits in visual acuity and motor function rather than a spatial learning and memory impairment. Thus, a visible (cued) platform version of the MWT was used in experiment 2 to rule out the involvement of sensory, motor and motivation deficits as non-cognitive confounding factors during spatial navigation within the task. We did not use the visible platform test in experiment 1 because it was a preliminary experiment. Moreover, we did not complete any experimental interventions in the C57 mice during experiment 1 that could potentially impair visual acuity, motor function, and/or motivation. The same pool and room were used for testing, with the exception that the platform was made visible by keeping the platform 1-2 cm above the water level and marked with black tape so that mice could locate the platform using the local visual cues rather than relying on extra-maze cues. The cued-platform task was carried out for 2 days after performing the competition test and consisted of one training session each day. Each session contained 8 trials (lasting 60 s) using each start point (N, S, E, W) once. The platform was placed at different locations during the sessions 1 and 2.

### Statistical analysis

The results are presented as mean±SEM. The results were analyzed using analysis of variance (ANOVA; SPSS 22, SPSS Inc., USA). One way ANOVA with repeated measures was used to detect differences between groups during phase one and two. The corridor percent path was analyzed using a paired samples *t*-test. Also, the paired samples *t*-test was used to measure differences between groups during the probe trial and competition test.

## Results

### Experiment 1

#### Phase one: original location training

As can be seen in figure 1B,C,G,H, mice showed a clear improvement in locating the hidden platform over the training days showing a typical learning curve in the standard version of the water task. This finding was confirmed statistically, and ANOVA indicated a statistically significant (F_(7,49)_ = 19.45. P = 0.000) decrease in latency to reach the hidden platform from day 1 (48.55 ± 2.49 sec) to day 8 (12.51 ± 1.55 sec, Fig. 1B). The distance mice travelled before reaching the platform also decreased over the training days (Fig. 1C) and this improvement in locating the platform was statistically significant across the training days. An analysis conducted on swim speed during spatial navigation in the acquisition phase did not show any statistically significant within-group differences indicating a normal motor and motivational function in the mice (Fig. 1E).

**Figure 1.**
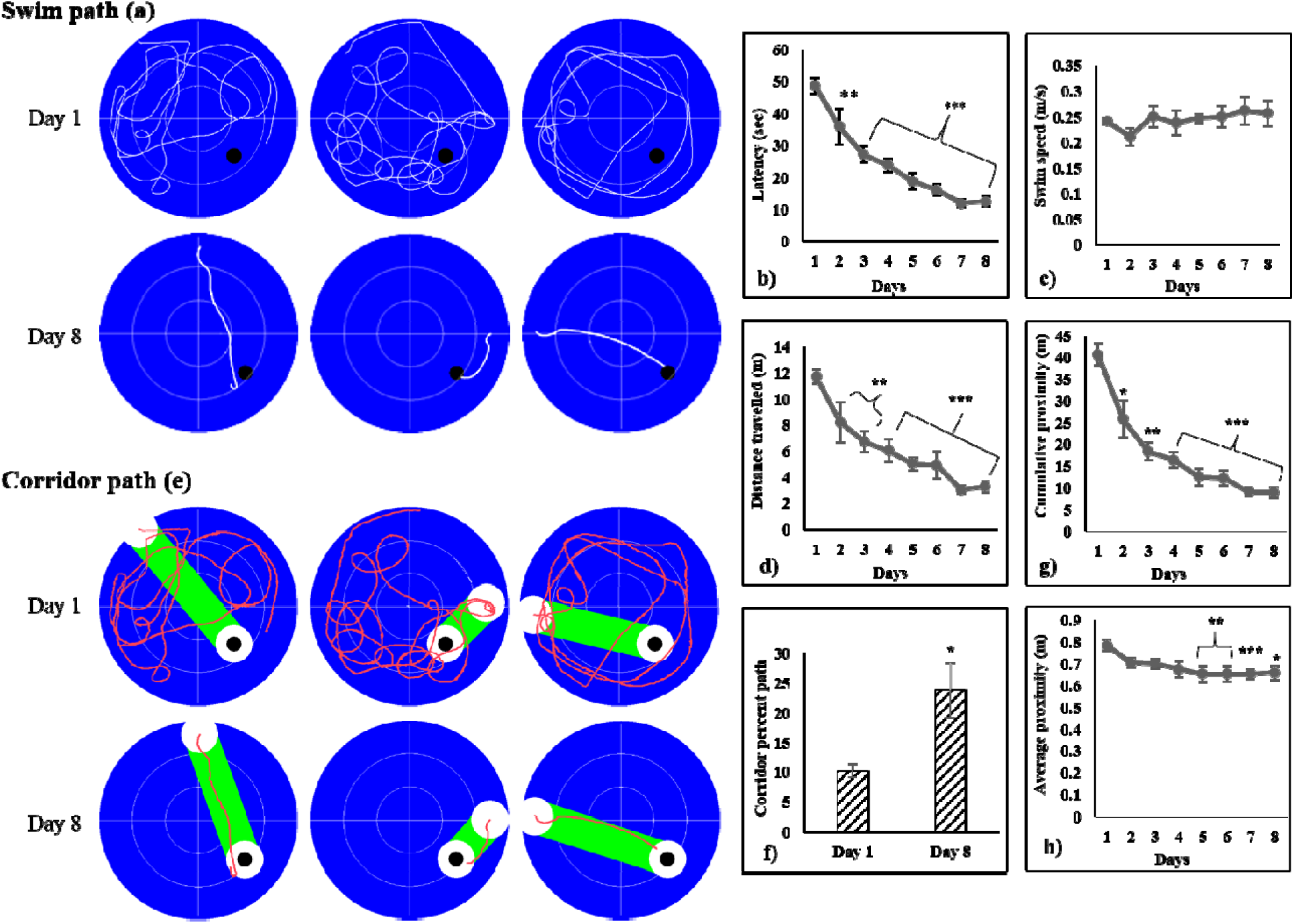
Evaluation of spatial learning performance of C57BL/6 mice during acquisition phase in Morris water task via analyzing A) swim path during the acquisition phase indicating the learning power of mice; B) latency to reach the platform; C) distance travelled by mice before reaching the platform; D) corridor percent path in the acquisition phase showing direct swim path to platform; E) swim speed during the acquisition phase; F) corridor percent path; G) cumulative proximity to platform and H) average proximity to platform. The results were expressed as mean±SEM. *P < 0.05; **P < 0.01; ***P < 0.001 as compared with day 1. The black spot in swim path and corridor path represents platform area.

Also, a paired *t*-test indicated that the corridor percent path was significantly (t_(7)_ = −3.381, P = 0.012) increased from 10.26 ± 1.04% on day 1 to 23.68 ± 4.67% on day 8 (Fig. 1F). Repeated measure ANOVA also revealed a statistically significant difference in Gallagher’s measures such as cumulative (F_(7,49)_ = 22.540, P = 0.000) and average (F_(7,49)_ = 2.307, P = 0.041) proximities to platform during the training (Fig. 1G,H).

#### Probe trial

The results from the probe trial showed a strong preference for the previously reinforced quadrant location as indicated by significantly greater search time in the training compared with other quadrants of the pool for both the 30 and 60 sec probe analyses (Fig. 2B). Analysis of the first half of the probe trial (30 sec duration) revealed that mice spent significantly (t_(7)_ = 3.063, P = 0.018) more time (47.5 ± 7.36%) in the target quadrant to search the platform location when compared with average time spent in other quadrants (17.44 ± 2.44%). Percent time spent by mice in the target quadrant was also found significantly (t_(7)_ = 3.636, P = 0.008) greater in comparison to the average time in other quadrants in probe trial duration of 0-60 sec. Moreover, as seen in Figure 2C, the distance travelled in the target versus the other quadrants was higher. When we analyzed the percent distance travelled by mice in the target versus other quadrants for the first 30 sec probe trial (t_(7)_ = 8.380, P = 0.000) and 0-60 sec (t_(7)_ = 4.031, P = 0.005), paired t-test showed that percent distance travelled by mice in target quadrant was significantly more in comparison to the other quadrants (Fig. 2C).

**Figure 2.**
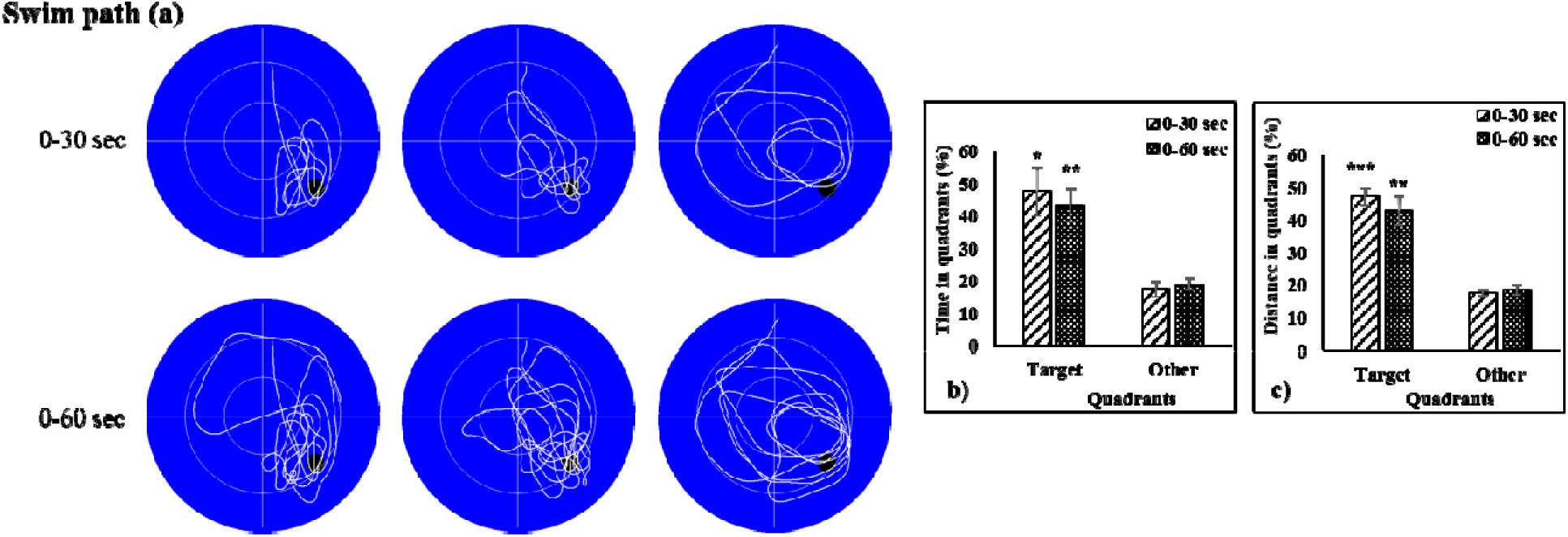
Evaluation of reference memory of C57BL/6 mice during probe trial in Morris water task via analyzing A) swim path indicating the reference memory of mice during 0-30 sec and 0-60 sec probe trials; B) time in target quadrant, and C) distance in target quadrant. The results were expressed as mean±SEM. *P < 0.05; **P < 0.01; ***P < 0.001 as compared with other quadrants. The black spot in swim path during probe trial of 0-30 sec & 0-60 sec represents platform area.

#### Phase two: new location training (rapid mass training)

In this phase, mice were trained for 20 trials, and data are presented in the form of trial blocks. Each trial block consisted of 4 trials from each starting point; therefore, animals had been trained in 5 trial blocks in this phase. The mice were able to rapidly learn a new spatial position in the same apparatus and testing room as original location training shown by decreases in latency and distance travelled to find the platform across training session (Fig. 3B,C). A statistical analysis of these measures of place learning showed that latency to reach the platform was decreased significantly (F_(4,28)_ = 4.410, P = 0.007) from 24.76 ± 4.89 sec in trial block 1 to 9.96 ± 1.39 sec in trial block 5. The distance travelled by mice to reach the platform was also significantly (F_(4,28)_ = 8.395, P = 0.000) shorten from 6.31 ± 0.93 m in trial block 1 to 2.00 ± 0.24 m in trial block 5. The swim speed of mice did not change significantly (F_(4,28)_ = 1.104, p = 0.374) from trial block 1 to trial block 5 (Fig. 3E). Paired *t*-test also showed a significant difference between trial blocks 1 and 5 in terms of the corridor percent path (t_(7)_ = −2.819, P = 0.026) suggesting that animals displayed greater spatial function in trial block 5 (27.53 ± 4.07%) than trial block 1 (14.30 ± 1.42%) in the mass training phase (Fig. 3f). An analysis conducted for shifting behavior from the old location quadrant (location during the acquisition phase) to the new location quadrant (opposite to the old quadrant location) indicated that mice significantly (t_(7)_ = 3.053, P = 0.018) preferred to search in the old location quadrant as compared with the new location quadrant in trial block 1 (Fig. 3G) indicating a robust memory for old location quadrant. Repeated measure ANOVA used to assess the trial block effect on mice learning also revealed that time spent in the quadrant with old location quadrant was decreased significantly (F_(4,28)_ = 7.936, P = 0.000) in trial block 5 as compared with trial block 1 (Fig. 3G). Similarly, statistically significant (F_(4,28)_ = 4.851, P = 0.004) increase in time spent in the new location quadrant was found in trial block 5 in comparison to trial block 1 (Fig. 3G). Moreover, the quadrant preference of mice was significantly (t_(7)_ = −2.487, P = 0.042) changed from the old location quadrant to new location quadrant in trial block 5 (Fig. 3G). A statistically significant difference in cumulative (F_(4,28)_ = 4.998, P = 0.004) and average (F_(4,28)_ = 6.684, P = 0.001) proximities to the platform was also found across the trial blocks (Fig. 3H,I).

**Figure 3.**
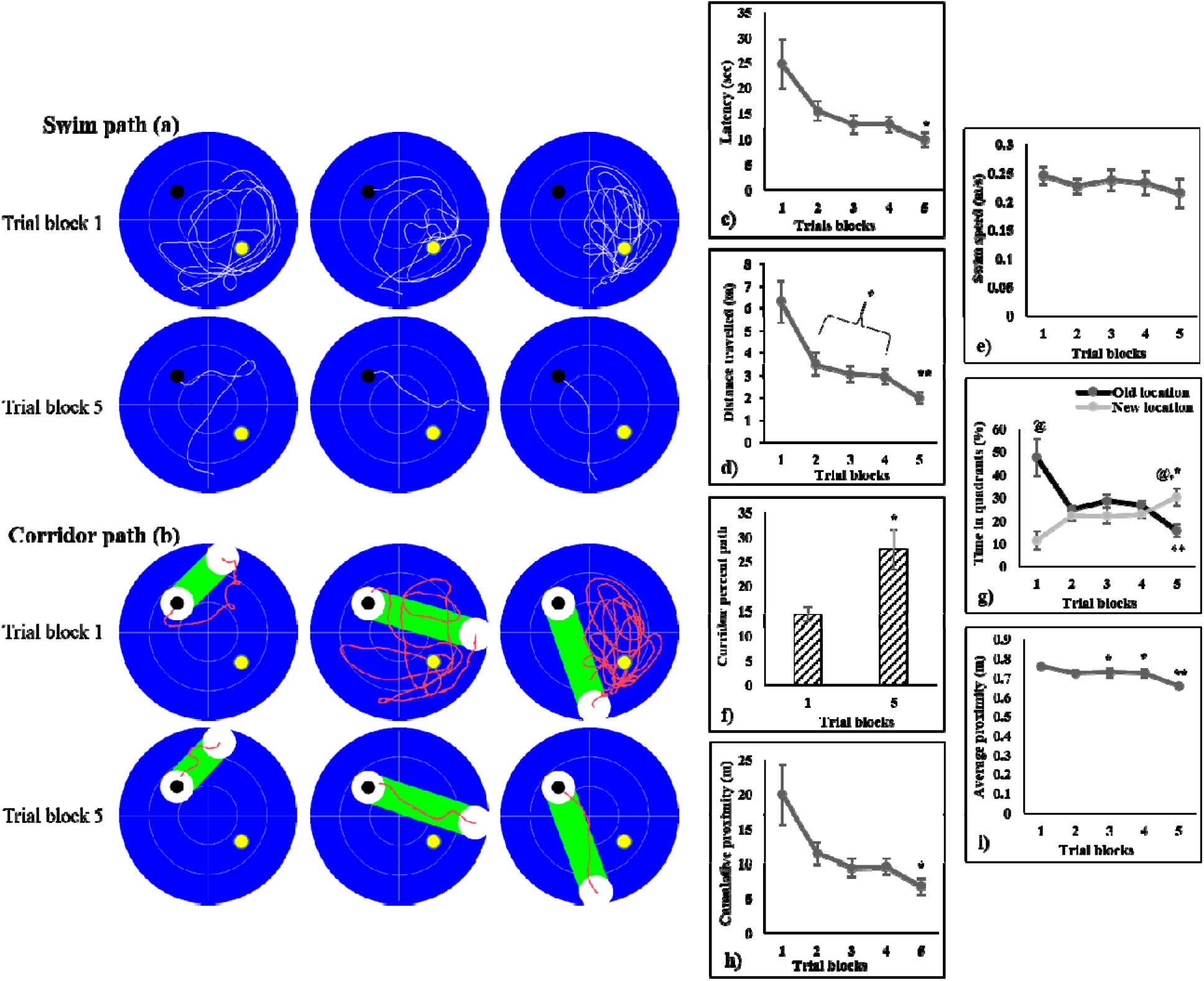
Evaluation of spatial learning performance of C57BL/6 mice during mass training phase in Morris water task via analyzing A) swim path during the mass training phase indicating the learning power of mice; B) latency to reach the platform; C) distance travelled by mice before reaching the platform; D) corridor percent path in the mass training phase showing direct swim path to platform; E) swim speed during the mass training phase; F) corridor percent path; G) time in quadrants with old and new location quadrants; H) cumulative proximity to platform, and I) average proximity to platform. The results were expressed as mean±SEM. *P < 0.05; ** P < 0.01; as compared with trial block 1. @P < 0.05 in between old and new location quadrants in trial block 1 & 5. The black and yellow spots in swim path and corridor path represent new & old quadrants area, respectively.

#### Phase three: competition test

The competitive probe trial provides an idea about the old memory retention and new memory formation in mice. The findings showed that mice spent more times in the old location quadrant indicating a preference for the old memory retention over the new memory formation. Analysis of the first half of the probe trial (0-30 sec) during the competitive probe trial indicated that mice spent significantly more times in the old location quadrant in comparison to the new location quadrant (t_(7)_ = 2.403, P = 0.047) as well as the other quadrants (t_(7)_ = 5.206, P = 0.001; Fig. 4B). Time spent by mice in the new location quadrant was also significantly (t_(7)_ = 3.227, P= 0.015) more as compared with the other quadrants in the first half of the probe trial (Fig. 4B). In the 60-sec probe trial, mice also spent more time (32.57 ± 4.19%) in the old location quadrant in comparison to the new location quadrant (25.66 ± 3.57%), though this difference was statistically non-significant (t_(7)_ = 0.907, P = 0.395; Fig. 4B). In comparison to the other quadrants, mice also spent significantly more time in the old (t_(7)_ = 2.607, P = 0.035) as well as the new (t_(7)_ = 2.430, P = 0.045) location quadrants during the 60-sec probe trial (Fig. 4B).

**Figure 4.**
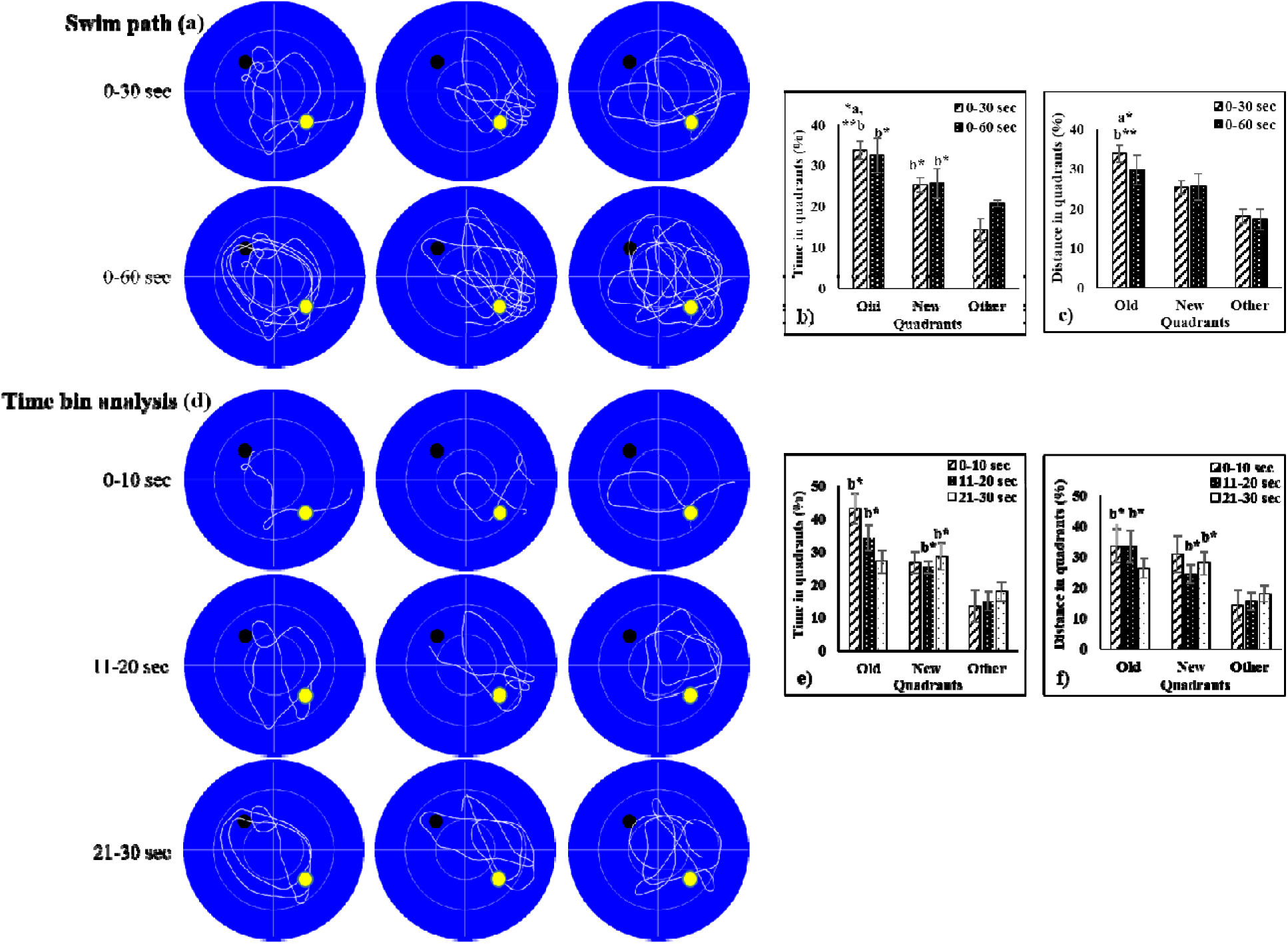
Evaluation of reference memory of C57BL/6 mice during competitive probe trial in Morris water task via analyzing A) swim path indicating the reference memory of mice during probe trial; B) time in target quadrant during 0-30 sec & 0-60 sec probe trials; C) distance in target quadrant during probe trial 0-30 sec & 0-60 sec. D) swim path during time bin analysis; E) time in target quadrant during probe trial 0-10, 11-20 & 21-30 sec, and F) distance in target quadrant during probe trial 0-10, 11-20 & 21-30 sec. The results were expressed as mean±SEM. *P < 0.05; **P < 0.01, a-as compared with new quadrant; b-as compared with other quadrant. The black and yellow spots in swim path represent new & old quadrants area, respectively.

Also, examination of the quadrant preference in terms of the distance travelled during the first half of probe trial (0-30 sec) revealed that distance travelled by mice was significantly more in the old location quadrant in comparison to the new location quadrant (t_(7)_ = 2.403, P = 0.047) and other quadrants (t_(7)_ = 5.829, P = 0.001) whereas the difference between percent distance travelled in the quadrant with the new location and other quadrants were close to a statistically significant level (t_(7)_ = 2.277, P = 0.057). However, no statistically significant difference was found in the distance travelled by mice in between the quadrants during the probe of 0-60 sec (Fig. 4C).

Additionally, the probe trial was analyzed across 10-sec time bins, and results indicated that mice spent more time in the old location quadrant in the first two 10-sec time bins showing a preference for the old versus the new locations. We found that mice spent more time in the old location quadrant in comparison to the new location quadrant and the other quadrants during the first (0-10 sec) and second (11-20 sec) time bins; however, a statistically significant difference was found only in comparison to the other quadrants (P < 0.05) as shown in figure 4E. In the third time bin (21-30 sec), mice spent significantly (P < 0.05) more time in the new location quadrant in comparison to the other quadrants. However, no statistically significant difference was found in the time spent by mice in the old location quadrant and the other quadrants (Fig. 4E). Similar results were found for distance travelled by mice in different quadrants in the third 10-sec interval (Fig. 4F). The results from this time bins analysis revealed that, under these training conditions, an older spatial memory representation dominated choice behavior over a new memory.

## Experiment 2

### Phase one: original location training

Similar to experiment 1, C57 mice showed a typical learning curve for the standard version of the MWT which is indicated by a decrease in escape latency to find the hidden platform over the training days whereas this learning pattern was lacking in *App*^*NL-G-F/NL-G-F*^ mice (Fig. 5B,C,G,H). Examination of acquisition in terms of the time spent to locate the hidden platform revealed a statistically significant (F_(7,84)_ = 21.518, P = 0.000) decrease in latency from day 1 (42.14 ± 3.64 sec) to day 8 (12.26 ± 0.96 sec) indicating that C57 animals took progressively shorter search times to find the platform as training proceeded over 8 days of testing (Fig. 5B). Moreover, C57 mice travelled significantly (F_(7,84)_ = 11.930, P = 0.000) less distance to find the hidden platform on day 8 (2.74 ± 0.22 m) than day 1 (7.66 ± 0.72 m) which clearly shows the learning ability of this mice strain (Fig. 5C). These mice were able to acquire and retrieve the spatial information over all training days compared with day 1 (Fig. 5C). However, *App*^*NL-G-F/NL-G-F*^ mice did not show any statistically significant difference in the latency (F_(7,49)_ = 1.358, P = 0.244) and distance travelled (F_(7,49)_ = 0.605, P = 0.749) on day 1 and 8 (Fig. 5B,C) which clearly indicates a deficit in learning ability of these knock-in mice. Moreover, *App*^*NL-G-F/NL-G-F*^ mice took significantly more time to reach the platform (t_(19)_ = −3.127, P = 0.006) and travelled more distance (t_(19)_ = −3.223, P = 0.004) on day 8 when compared with C57 mice indicating a difference in the *App*^*NL-G-F/NL-G-F*^ mice and controls in their ability to acquire the spatial information (Figs. 5B,C). Examination of swim speed during acquisition showed that animals swam somewhat faster on days 5-7. However, no statistically significant differences were observed between days (Fig. 5E).

**Figure 5.**
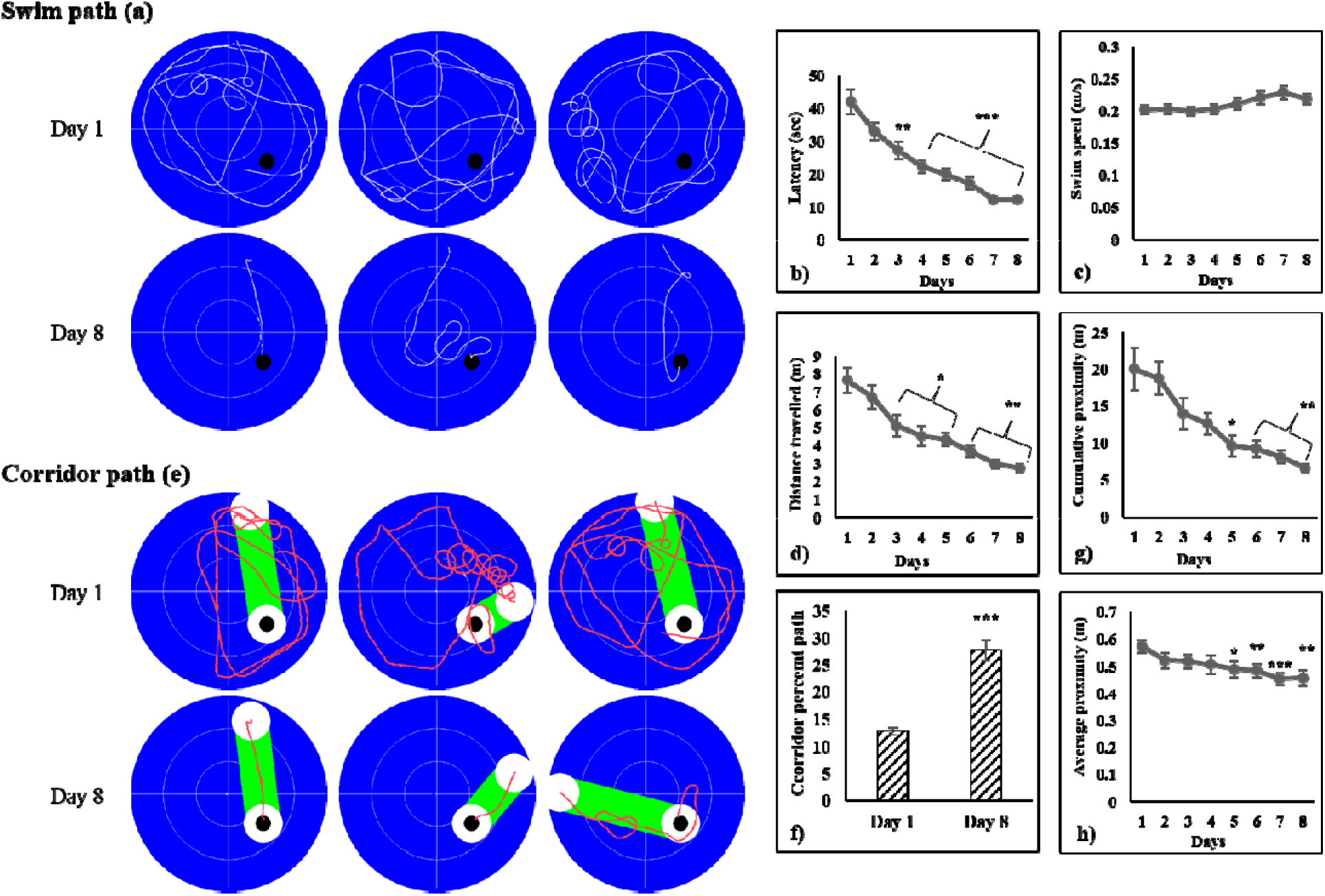
Evaluation of spatial learning performance of C57BL/6 and AppNL-G-F/NL-G-F (ADTg) mice during acquisition phase in Morris water task via analyzing A) swim path during the acquisition phase indicating the learning power of mice; B) latency to reach the platform; C) distance travelled by mice before reaching the platform; D) corridor percent path in the acquisition phase showing direct swim path to the platform; E) swim speed during the acquisition phase; F) corridor percent path; G) cumulative proximity to the platform and H) average proximity to the platform. The results were expressed as mean±SEM. *P < 0.05; **P < 0.01; ***P < 0.001, a-as compared with day 1 for C57 mice; b-as compared with AppNL-G-F/NL-G-F (ADTg) mice on day 8; c-as compared with C57 mice. The black spot in swim path and corridor path represents platform area.

Moreover, in C57 mice, corridor percent path was increased significantly (t_(12)_ = −7.523, P = 0.000) from 12.79 ± 0.72 on day 1 to 27.75 ± 1.91 on day 8 which shows a tendency to swim in a straight path towards the escape platform is increased on day 8 as compared with day 1 (Fig. 5F). Further evidence of improved spatial specificity in the C57 mice comes from and analysis of cumulative (F_(7,84)_ = 10.555, P = 0.000) and average proximities (F_(7,84)_ = 8.833, P = 0.000) to platform, measurements of Gallagher, were also significantly decreased across the training days (Figs. 5G,H) indicating that on day 8, C57 mice were spending more time close to platform. All the parameters measured during the acquisition phase showed that C57 mice learned the specific location of the hidden platform. In contrast, no statistically significant difference was found in corridor percent path (t_(7)_ = 0.207, P = 0.842), cumulative (F_(7,49)_ = 1.718, P = 0.127) and average (F_(7,49)_ = 1.321, P = 0.269) proximities to platform on day 8 in comparison to day 1 for *App*^*NL-G-F/NL-G-F*^ mice indicating an impairment in spatial learning (Figs 5F,G,H). Furthermore, *App*^*NL-G F/NL-G-F -*^ mice showed less corridor percent path (t_(19)_ = 3.700, P = 0.002), higher cumulative (t_(19)_= −2.936, P = 0.008) and average proximities (t_(19)_ = −2.391, P = 0.027) on day 8 as compared with C57 mice (Figs 5F,G,H). This pattern of results indicate that *App*^*NL-G-F/NL-G-F*^ mice showed an impairment of spatial learning ability.

#### Probe trial

The results from the probe trial indicated a strong spatial reference memory in C57 mice which is indicated by significantly more time spent by mice in the target quadrant compared with other quadrants of the task for both the 30- and 60-sec analyses (Fig. 6B). In the probe trial of 0-30 sec, percent time spent in target quadrant by C57 mice was significantly (t_(12)_= 4.878, P = 0.000) higher in comparison to average percent time spent in other quadrants suggesting that animals exhibited a less diffuse pattern of searching, with much more spatial bias toward the former training quadrant (Fig. 6B). Similar results were found for 0-60 sec probe trial where C57 mice spent significantly (t_(12)_ = 6.324, P = 0.000) more time in target quadrant as compared with other quadrants (Fig. 6B). Paired *t*-test also showed a statistically significant difference in percent distance travelled by C57 mice in target quadrant and average of other quadrants during the probe trial of 0-30 sec (t_(12)_ = 5.098, P = 0.000) and 0-60 sec (t_(12)_ = 6.610, P = 0.000). C57 mice travelled 40.58 ± 3.07% in the target quadrant in comparison to 19.74 ± 1.02% in average of other quadrants in probe trial 0-30 sec (Fig. 6C). Similarly, in probe trial of 0-60 sec, the distance travelled by C57 mice in the target quadrant (39.61 ± 2.24%) was significantly higher in comparison to average of other quadrants (19.49 ± 0.74%; Fig. 6C). However, no statistically significant difference was found in time spent (for 30 sec, t_(7)_ = 0.381, P = 0.715; for 60 sec, t_(7)_ = 1.720, P = 0.129) and distance travelled (for 30 sec, t_(7)_ = 0.378, P = 0.717; for 60 sec, t_(7)_ = 1.770, P = 0.120) by *App*^*NL-G-F/NL-G-F*^ mice in target quadrant as compared with the average of other quadrants (Figs 6B,C). Furthermore, *App*^*NL-G-F/NL-G-F*^ mice significantly spent less time (for 30 sec, t_(19)_ = 3.041, P = 0.007; for 60 sec, t_(19)_ = 2.346, P = 0.030) and travelled less distance (for 30 sec, t_(19)_ = 3.236, P = 0.004; for 60 sec, t_(19)_ = 2.616, P = 0.017) in the target quadrant as compared with C57 mice for 30-sec and 60-sec probe trial indicating an impairment in spatial memory (Fig. 6B,C).

**Figure 6.**
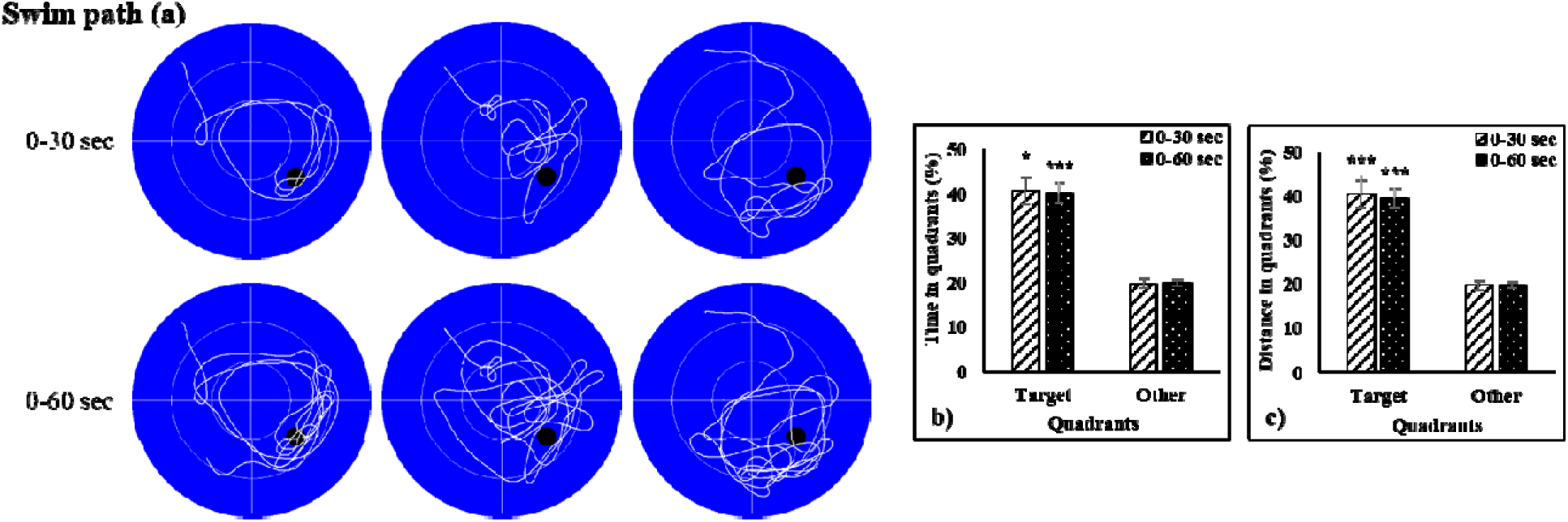
Evaluation of reference memory of C57BL/6 and AppNL-G-F/NL-G-F (ADTg) mice during probe trial in Morris water task via analyzing A) swim path indicating the reference memory of mice during 0-30 sec and 0-60 sec probe trials; B) time in target quadrant, and C) distance in target quadrant. The results were expressed as mean±SEM. *P < 0.05; **P < 0.01; ***P < 0.001, a-as compared with other quadrants; b-as compared with AppNL-G-F/NL-G-F (ADTg) for the target quadrant. The black spot in swim path during probe trial of 0-30 sec & 0-60 sec represents platform area.

#### Phase two: new location training (rapid mass training)

During this phase, 32 trials were given to each mouse and data are presented in the form of 8 trial blocks each consisting of 4 trials. Like the results from experiment 1, decreases in latency and distance travelled by mice to find the platform across training trials were observed showing the ability of mice to rapidly learn a new spatial position in the same apparatus and testing room as the original location. The latency to reach the platform was decreased significantly (F_(7,84)_ = 14.112, P = 0.000) from 35.78 ± 4.64 sec in trial block 1 to 12.06 ± 0.911 sec in trial block 8 (Fig. 7B) showing that the animals were able to locate the hidden platform. The distance travelled by mice to reach the platform was also significantly (F_(7,84)_= 10.083, P = 0.000) decreased from 6.39 ± 0.80 m in trial bock 1 to 2.15 ± 0.29 m in trial block 8 (Fig. 7C). Moreover, in this phase, *App*^*NL-G-F/NL-G-F*^ mice also showed a learning behaviour which was indicated by statistically significant decreases in latency (t_(7,49)_ = 7.191, P = 0.000) and distance travelled (t_(7,49)_ = 8.402, P = 0.000) in trial block 8 as compared with trial block 1 (Figs 7B,C). However, *App*^*NL-G-F/NL-G-F*^ mice took significantly more time to14 reach the platform (t_(19)_ = −2.102, P = 0.049) and travelled more distance (t_(19)_ = −2.156, P = 0.044) in trial block 8 when compared with C57 mice indicating that *App*^*NL-G-F/NL-G-F*^ mice are not as good as C57 mice to learn during this phase of training (Figs 7B,C). The swim speed of mice in both groups was not changed significantly throughout rapid mass training phase (Fig. 7D). Moreover, the corridor percent path, for C57 mice was increased significantly (t_(12)_= −6.187, P = 0.000) from trial block 1 (14.89 ± 2.00%) to trial block 8 (26.68 ± 2.35%) indicating an increase in spatial accuracy during training (Fig. 7E). However, in *App*^*NL-G-F/NL-G-F*^ mice, no statistically significant (t_(7)_ = −2.169, P = 0.067) difference was found between the corridor percent path in trial block 1 and trial block 8. An analysis conducted for shifting pattern of search strategy of C57 and *App*^*NL-G-F/NL-G-F*^ mice from the old location quadrant (location during the acquisition phase) to the new location quadrant (opposite to the old location quadrant) indicated that the quadrant preference of C57 and *App*^*NL-G-F/NL-G-F*^ mice was significantly (t_(12)_ = 7.328, P = 0.000 for C57; t_(6)_ = 2.671, P = 0.037 for *App*^*NL-G-F/NL-G-F*^) higher for old location quadrant as compared with new location quadrant in trial block 1 indicating a memory for old location quadrant (Fig. 7F). However, in trial block 8, these mice showed significantly (t_(12)_ = −5.566, P = 0.000 for C57; t_(12)_ = −5.155, P = 0.001 for *App*^*NL-G-F/NL-G-F*^) more preference for the new location quadrant as compared with old location quadrant indicating that both groups of mice can learn the new location quadrant (Fig. 7F). A repeated measure ANOVA was also used to assess the trial block effect on the learning ability of mice. Mice spent significantly (F_(7,84)_= 21.693, P = 0.000 for C57; F_(7,49)_ = 3.677, P = 0.003 for *App*^*NL-G-F/NL-G-F*^) less time in the old location quadrant in trial block 8 when compared with trial block 1. Also animals spent significantly (F_(7,84)_= 10.922, P = 0.000 for C57; F_(7,49)_ = 7.526, P = 0.000 for *App*^*NL-G-F/NL-G-F*^) more time in the new location quadrant on trial block 8 as compared with trial block 1 (Fig. 7F). However, no statistically significant difference was found between C57 and *App*^*NL-G-F/NL-G-F*^ mice in the time spent in new location quadrant in trial block 8 (Fig. 7F).

**Figure 7.**
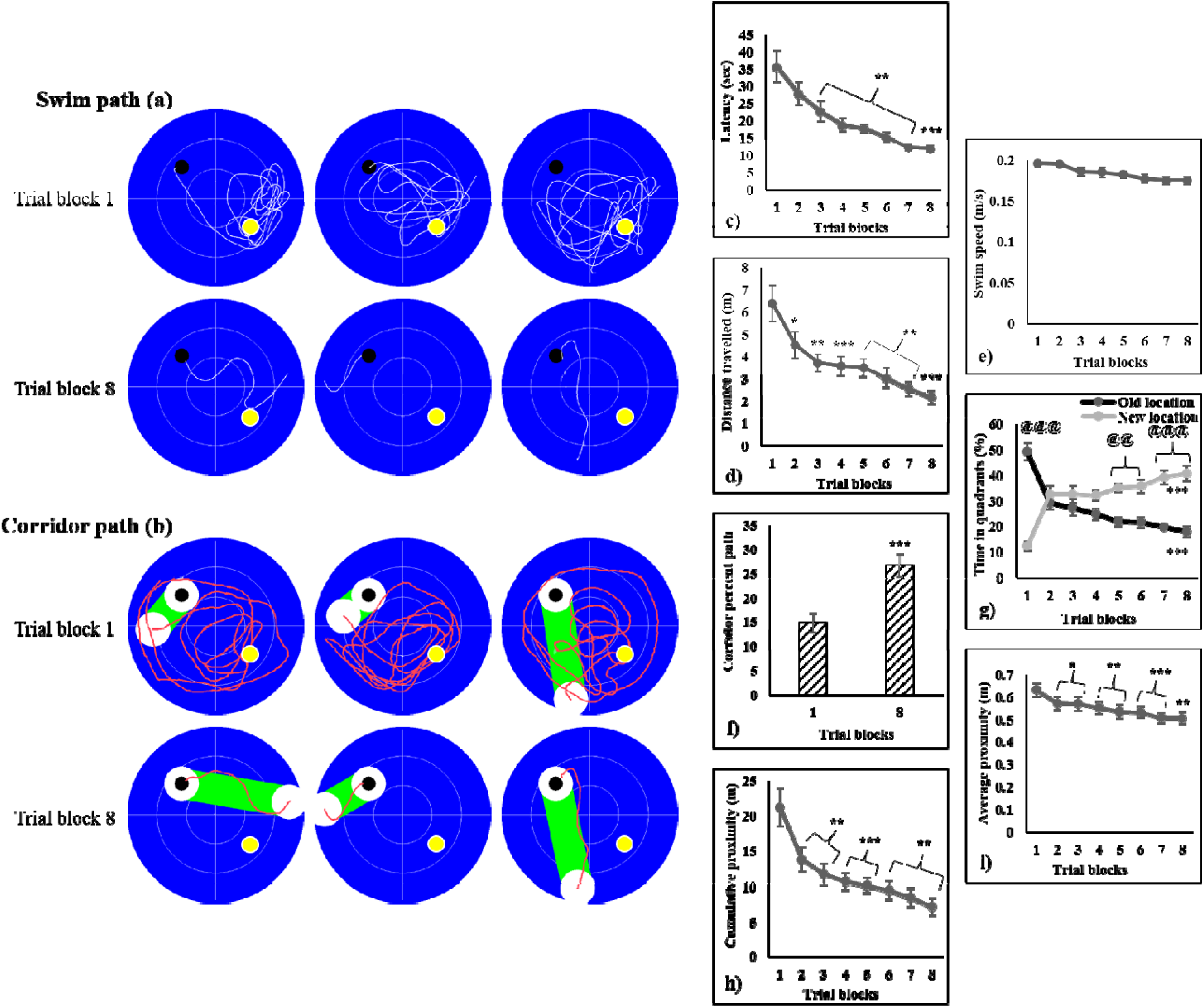
Evaluation of spatial learning performance of C57BL/6 and AppNL-G-F/NL-G-F (ADTg) mice during mass training phase in Morris water task via analyzing A) swim path during the mass training phase indicating clear rapid spatial learning in mice; B) latency to reach the platform; C) distance travelled by mice before reaching the platform; D) swim speed during the mass training day; E) corridor percent path; F) shifting pattern from quadrant with old location quadrant to new location quadrant; G) corridor percent path in the mass training phase showing direct swim path to platform; H) cumulative proximity to platform, and I) average proximity to platform. The results were expressed as mean±SEM. *P < 0.05; **P < 0.01; ***P < 0.001, a-as compared with trial block 1 for C57 mice; b-as compared with trial block 1 for AppNL-G-F/NL-G-F (ADTg) mice; c-as compared with C57 mice; d-as compared with the old location quadrant for C57 mice; e-as compared with old location quadrant for AppNL-G-F/NL-G-F (ADTg) mice. $$$P < 0.001 as compared with the new location quadrant for C57 mice; #P < 0.05 as compared with the new location quadrant for AppNL-G-F/NL-G-F (ADTg) mice. The black and yellow spots in swim path and corridor path represent the new and old quadrants area, respectively.

Also, repeated measure ANOVA showed a statistically significant improvement in cumulative (F_(7,84)_= 10.555, P = 0.000 for C57; F_(7,49)_= 7.285, P = 0.000 for *App*^*NL-G-F/NL-G-F*^) and average proximities (F_(7,84)_ = 8.833, P = 0.000 for C57; F_(7,49)_= 7.430, P = 0.000 for *App*^*NL-G-F/NL-G-F*^) to the platform location during the training sessions (Fig. 7H,I). However, no statistically significant difference was found in corridor percent path, cumulative and average proximities in trial block 8 in between C57 and *App*^*NL-G-F/NL-G-F*^ mice (Figs. 7E,H,I). The results from the mass training phase showed learning ability of both C57 and *App*^*NL-G-F/NL-G-F*^ mice.

#### Phase three: competition test

Using the competitive probe trial, we determined the preference for the old versus new location quadrants in mice. The findings showed that C57 mice spent approximately the same amount of time and travelled the same distance in both target quadrants (old and new) indicating an equal preference for the old and new escape locations. In a bin analysis of the competitive probe trial at 0-30 sec, it was found that C57 mice spent significantly more time in the old (t_(12)_ = 2.625, P = 0.022) and new location quadrants (t_(12)_= 3.214, P = 0.007) in comparison to the other quadrants (Fig. 8B). Similarly, a bin analysis of the 0-60 sec probe trial showed that C57 mice spent significantly more time in the old (t_(12)_= 2.543, P = 0.026) and new location quadrants (t_(12)_= 3.588, P = 0.004) as compared with other quadrants (Fig. 8B). However, *App*^*NL-G-F/NL-G-F*^ mice showed a different pattern. They spent significantly more time in the new location quadrant as compared with the old location quadrant and the other quadrants for 30 sec (t_(7)_= 3.315, P = 0.013 new vs old; t_(7)_= 2.895, P = 0.023 new vs other) and 60 sec (t_(7)_= 5.266, P = 0.001 new vs old; t_(7)_= 3.000, P = 0.020 new vs other) probe trials (Fig. 8B). Similarly, C57 mice travelled significantly more distance in the old location quadrant (t_(12)_= 2.395, P = 0.034) as well as the new location quadrant (t_(12)_= 2.544, P = 0.026) in comparison to the other quadrants in the probe trial of 0-30 sec (Fig. 8C). Moreover, distance travelled by mice in the old (t_(12)_= 2.372, P = 0.035) and new location quadrants (t_(12)_= 3.393, P =0.005) was significantly higher in comparison to other locations in the pool during the 0-60 sec probe trial (Fig. 8C). However, no statistically significant difference was found in the time spent as well as the distance travelled by C57 mice in the old and new location quadrants during the competitive probe trial of 0-30 sec and 0-60 sec (Figs. 8B-C). Furthermore, *App*^*NL-G-F/NL-G-F*^ mice travelled significantly more distance in the new location quadrant when compared with the old location quadrant and other quadrants for 30 sec (t_(7)_= 3.405, P = 0.011 new vs old; t_(7)_= 3.054, P = 0.018 new vs other) and 60 sec (t_(7)_= 4.309, P = 0.004 new vs old; t_(7)_= 2.987, P = 0.020 new vs other) probe trials (Figs 8B,C). However, no statistically significant difference was found in time spent as well as distance travelled by *App*^*NL-G-F/NL-G-F*^ mice between the old location quadrant versus the other quadrants during the competitive probe trial of 0-30 sec and 0-60 sec (Figs. 8B-C). Additionally, *App*^*NL-G-F/NL-G-F*^ mice spent significantly less time and travelled less distance in the old location quadrant in 30-sec (t_(19)_= 2.800, P = 0.011 for time spent; t_(19)_= 2.814, P = 0.011 for distance travelled) and 60-sec (t_(19)_= 2.954, P = 0.008 for time spent; t_(19)_= 2.308, P = 0.032 for distance travelled) probe trials when compared with C57 mice. This pattern of results indicates that *App*^*NL-G-F/NL-G-F*^ mice did not display retention of spatial memory for the old platform location. These results suggest that *App*^*NL-G-F/NL-G-F*^ showed a different pattern of behavior on the competition test compared to C57 mice. C57 mice showed evidence for retention of both the new and old locations during this test. In contrast, *App*^*NL-G-F/NL-G-F*^ mice only retained the new location memory and showed no evidence of the older memory.

**Figure 8.**
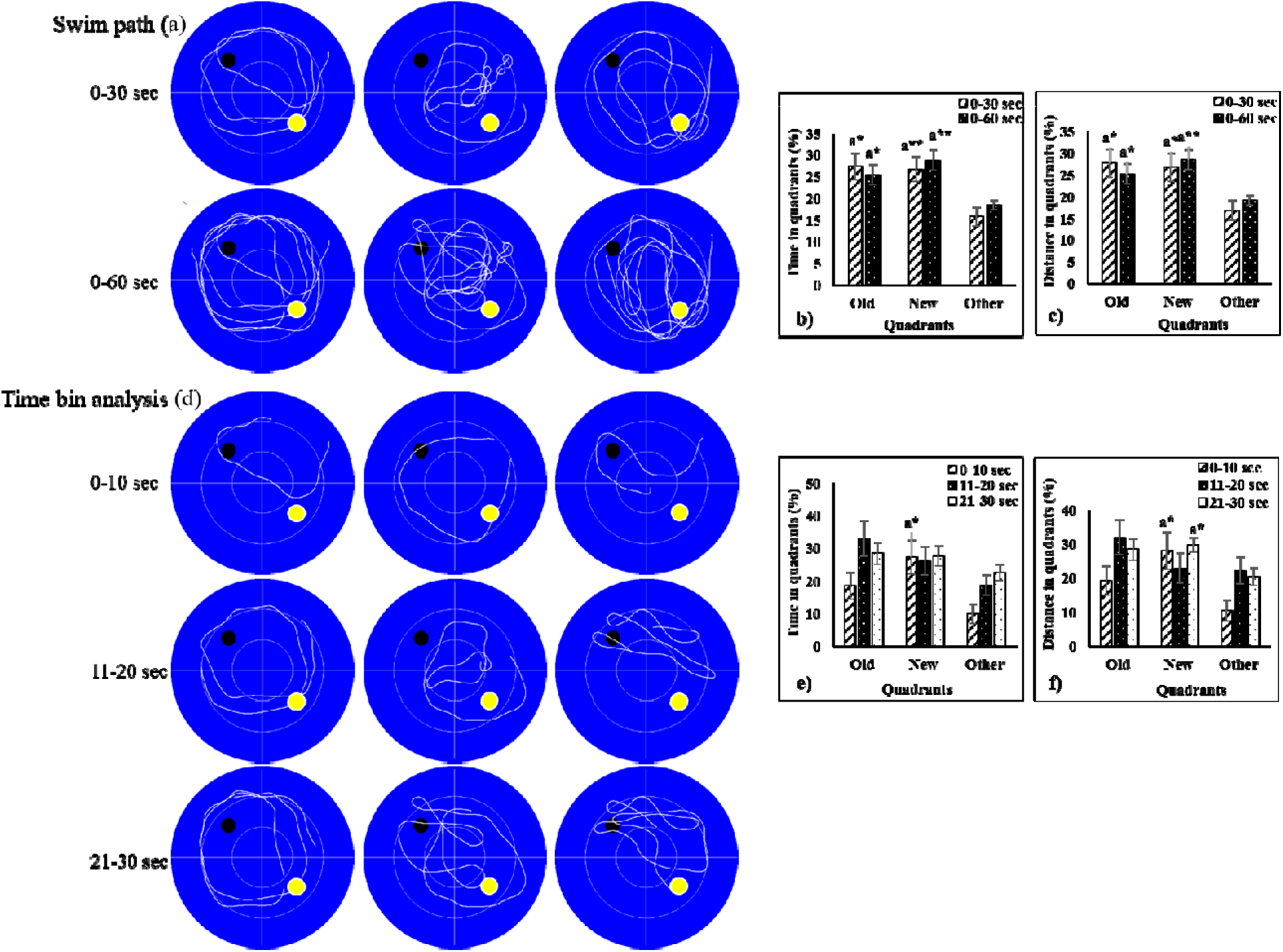
Evaluation of reference memory of C57BL/6 and AppNL-G-F/NL-G-F (ADTg) mice during competitive probe trial in Morris water task via analyzing A) swim path indicating the reference memory of mice during probe trial; B) time in quadrants during probe trial 0-30 sec & 0-60 sec; C) distance in quadrants during 0-30 sec and 0-60 sec probe trials; D) swim path during time bin analysis; E) time in quadrants during probe trial 0-10, 11-20 & 21-30 sec, and F) distance in quadrants during 0-10, 11-20 and 21-30 sec probe trials. The results were expressed as mean±SEM. *P < 0.05; **P < 0.01, a-as compared with other quadrant; b-as compared with AppNL-G-F/NL-G-F (ADTg) mice for old location quadrant; c-as compared with the old location quadrant for AppNL-G-F/NL-G-F (ADTg) mice; d-as compared with other quadrants for AppNL-G-F/NL-G-F (ADTg). The black and yellow spots in swim path represent the new and old quadrants area, respectively.

In a time bin analysis, it was found that C57 mice spent more time and travelled more distance in the old and new location quadrants compared to the other quadrants in the first 10 sec, however, a statistically significant difference was found only when comparing the new location quadrant to the other quadrants (P < 0.05) (Figs. 8E-F). During the 11-20-sec time bin, C57 mice spent more time in the old and new location quadrants compared to the other quadrants although this difference was not statistically significant (Fig. 8E) nor was the distance travelled by these mice in these quadrants. Similarly, in the 21-30 sec time bin, C57 mice spent approximately equal time in both the old and new location quadrants but the time spent by these mice in these target quadrants was more in comparison to the other quadrants (Fig. 8F). These mice travelled more distance in the new location quadrant as compared with the other quadrants (P < 0.05) but no statistically significant difference was found between the old location quadrant and other quadrants (Fig. 8F). *App*^*NL-G-F/NL-G-F*^ mice spent significantly more time (P < 0.01 new vs old; P < 0.05 new vs other) and travelled more distance (P < 0.05 new vs old; P < 0.05 new vs other) in the new location quadrant in comparison to the old location quadrant and the other quadrants for time bin 0-10 sec interval. For the 11-20 second time bin, a difference was found in time spent (P < 0.05) and distance travelled (P < 0.05) by *App*^*NL-G-F/NL-G-F*^ mice in between old and new location quadrants; however, no statistically significant difference was found between old and other quadrants, and new and other quadrants. Furthermore, *App*^*NL-G-F/NL-G-F*^ mice did not show any differences between the quadrants for time bin 21-30 sec. Likewise, *App*^*NL-G-F/NL-G-F*^ mice significantly spent less time (P < 0.01 for 0-10 sec; P <0.05 for 11-20 sec) and travelled less distance (P < 0.05 for 0-10 sec; P <0.05 for 11-20 sec) in the old location quadrant when compared with C57 mice for time bin 0-10 sec and 11-20 sec whereas no difference was found in the time spent and distance travelled in the new location quadrant. Additionally, no statistically significant difference was found between C57 and *App*^*NL-G-F/NL-G-F*^ mice in the time spent and the distance travelled in quadrants for time bin 21-30 sec. Moreover, an analysis of what quadrant the mice first entered for the first 10 sec time bin showed that 53.84% C57 mice opted to enter the old location quadrant and 46.16% mice entered into the new location quadrant, showing equal preference for old and new location quadrants whereas only 12.5% *App*^*NL-G-F/NL-G-F*^ mice opted to enter the old location quadrant and 87.5% mice opted to enter the new location quadrant. Taken together, the results from the time bin analysis during the competition probe trials revealed further evidence to suggest that C57 mice showed memory retention of both the old and new target locations whereas *App*^*NL-G-F/NL-G-F*^ mice seem to have no memory for the old target location or were unable to retrieve it.

*Visible platform test.* All animals in experiment 2 were tested in the cued version of the MWT that does not require spatial learning and memory functions. This task was also selected to rule out the possibility of the impact of non-cognitive factors on spatial navigation. The latency to reach the visible platform for C57 was decreased significantly (t_(12)_= 12.905, P= 0.000) from 13.62±0.55 sec in session 1 to 5.87±0.39 sec in session 2 (Fig. 9). Additionally, *App*^*NL-G-F/NL-G-F*^ mice also showed a decrease in latency to reach the platform from 14.78±0.87 sec in day 1 to 6.78±0.64 sec in day 2 (Fig. 9). Moreover, no statistically significant difference was found in latency between C57 and *App*^*NL-G-F/NL-G-F*^ mice. These results indicate that impairment in spatial memory found in *App*^*NL-G-F/NL-G-F*^ mice was not due to impairments in visual acuity, sensorimotor functions, or motivation.

**Figure 9:** Evaluation of non-spatial performance of C57BL/6 and AppNL-G-F/NL-G-F (ADTg) mice in the visible platform version of the Morris water task. The results were expressed as mean±SEM. ***P < 0.001 as compared with session 1.

## Discussion

In the present study, a protocol for MWT was optimized to assess old and new spatial memory of mice. All measures of spatial learning and memory function assessed in both experiments indicated robust spatial learning across multiple days of acquisition to a static platform location and clear memory for that location during a probe test. The mice also could rapidly learn and remember a new escape location in the same task and testing room in a single day of massed training. Interestingly, a competition test at the end of training in which the subjects were free to swim wherever they wanted showed that the mice had both new and old memories available to them. Overall, these results suggest that like rats, mice can learn multiple spatial locations in the MWT. In the following sections, we will discuss, in more detail, the pattern of results exhibited by mice during the different phases of training on this variant of the MWT.

### Original location training

In both sets of experiments, we found very consistent results during the original location phase and associated probe trial. As original location training unfolded, there was statistically significant decreases in latency as well as distance travelled over the training days with no statistically significant change in their swim speed throughout. The increased accuracy of the swim paths of C57 mice in experiment 1 & 2 also support the latency and distance travel data during the acquisition phase so that by day 8 of training in both the experiments the C57 mice are traveling straight to the platform indicating that mice can not only show improved spatial knowledge over training days, but they also show improvements in the precision of their spatial representation and related navigational abilities. Further evidence for the latter claim comes from the corridor percent path analysis in which the C57 mice spent more time in the corridor percent path on later in training compared with early trials and the Gallagher measurements like cumulative proximity distance and average proximity distance were significantly lower later versus early in training which means that C57 mice spent more time close to platform showing an increased spatial knowledge as training progressed. However, the analysis of various dependent variables in experiment 2 indicated that *App*^*NL-G-F/NL-G-F*^ mice did not learn this task displaying a severe impairment in cognitive function in *App*^*NL-G-F/NL-G-F*^ mice. Although in previous studies, average proximity to platform has been used to assess spatial memory of rats and mice during the probe trial^28,29^ results of the present study suggest that cumulative and average proximities to platform can also be used as an indicator of learning ability of rats and mice in the acquisition phase. Su et al. also used cumulative proximity to assess the learning behavior of mice and reported that as the training progresses, cumulative proximity decreases which support the findings of our study^28^

### Probe Trial: Original Location

In the probe trials of experiment 1 and 2, C57 mice spent significantly more time and travelled more distance in the target quadrant in comparison to other quadrants indicating a clear spatial memory for the correct escape location in these subjects. This conclusion is also supported by the pattern of swim paths exhibited by the mice during the probe trial of both experiments. In both experiments, C57 mice showed similar swim path pattern indicating more time spent by mice to locate the platform in the target quadrant (Figs. 2A, 6A). The results from the probe trial in the present study are in accordance with findings from 19 previous studies^30-32^. In contrast, *App*^*NL-G-F/NL-G-F*^ mice in experiment 2 did not show any statistically significant difference in time spent, and distance travelled in the target quadrant when compared with other quadrants. Moreover, these mice significantly spent less time and travelled less distance in the target quadrant in comparison to C57 mice. These results indicate a clear impairment of spatial memory retention in *App*^*NL-G-F/NL-G-F*^ mice.

### New location training (rapid mass training)

The learning pattern shown by C57 mice during the rapid new location learning phase in the present study is in agreement with findings of our previous work using rats^19^. During the new location training phase, C57 mice learned the new location of the platform in both experiments. An important demonstration in the current experiments was that the mice clearly showed the shifting pattern from old location quadrant to new location quadrant (Fig. 3G in experiment 1; Fig. 7F in experiment 2) which indicates that mice were learning the new location of the platform. Our analysis also indicates when the switch in preference for the two locations occurs, an effect that has never been demonstrated before in the mouse or rat. By the end of training in both experiments, mice completely shifted their quadrant preference from old to new. *App*^*NL-G-F/NL-G-F*^ mice also showed the ability to rapidly acquire a new platform location as indicated by all of the dependent variables except for corridor percent path. The latter finding indicating that these mice do not travel straight to platform position as C57 indicating a difference in their platform search strategies and spatial specificity of the representation acquired.

Moreover, a statistically significant difference was found in latency and distance travelled on trial block 8 between C57 and *App*^*NL-G-F/NL-G-F*^ mice indicating AD mice are not as good as C57 to learn this task. Quadrant preference in C57 mice for old location was also decreased significantly on trial block 5 & 8 as compared with trial block 1 in experiment 1 & 2, respectively (Fig. 3G in experiment 1; Fig. 7F in experiment 2). Similarly, C57 mice spent significantly more time in the new location quadrant in trial block 5 & 8 as compared with trial block 1 in experiment 1 & 2, respectively indicated that C57 mice easily learned the new location quadrant (Fig. 3G in experiment 1; Fig. 7F in experiment 2). Increased spatial accuracy of C57 mice was also evident by an increased corridor percent path on trial block 5 in experiment 1 (Fig. 3F) and trial block 8 in experiment 2 (Fig. 7E) as compared with trial block 1. Interestingly, *App*^*NL-G-F/NL-G-F*^ mice also showed a similar pattern of quadrant preference shifting from old to new memory. In the case of C57 mice, statistically significant quadrant preference shifting pattern was found from trial block 5-8, whereas, for *App*^*NL-G-F/NL-G-F*^ mice, this pattern was observed on trial block 7 & 8 only. This indicates that it took longer for *App*^*NL-G-F/NL-G-F*^ animals to shift to the new location quadrant compared to C57 mice. The corridor percent path analysis has been studied in rats^25^ and our findings are consistent with that study. In the case of mice, there is only one study^27^ that assessed corridor percent path analysis though it did not assess this measure during a rapid mass training phase to learn a new platform location in a single training session. Along with the other parameters of learning assessed in this version of the MWT, cumulative and average proximities to the platform also decreased over training which also supports the idea that C57 mice learned the specific location of the new platform. *App*^*NL-G-F/NL-G-F*^ mice did show evidence that they could acquire a new spatial location representation under these mass training conditions conducted in one session although it did take them longer to switch to a new location preference.

### Competitive probe trial (old versus new memory)

In a previous study, it has been reported that mice trained during a rapid mass training phase spent more time in the target quadrant in the probe trial indicating new memory formation of mice^27,33^. However, in these studies, they did not report on time spent in old location quadrant, and the training occurred over many days not in a single training session. From the findings of the previous studies, it was unclear whether the mice have any traces of their old spatial memory acquired in that apparatus and context. Accordingly, in the present study, we performed a competitive probe trial to assess the preference of mice towards old versus new memories. In experiment 1, C57 mice showed a preference for the old location quadrant. Therefore, in experiment 2, we increased the number of trials up to 32 so that mice show equal memory preference for both old and new location quadrants. In this study, C57 mice showed retention of both old and new spatial memories. *App*^*NL-G-F/NL-G-F*^ mice showed a different pattern of results indicating a specific type of memory impairment. *App*^*NL-G-F/NL-G-F*^ mice showed a preference for the new spatial location which suggests that *App*^*NL-G-F/NL-G-F*^ mice only retained the recently learned information which may more rapidly decay. Therefore, the new MWT protocol reported in experiment 2 can be employed to assess the old and new memory formation and retention in mice. The results of the present report strongly suggest that this new optimized protocol would be valuable for the assessment of impairment in learning and memory functions in experimental models for memory impairments in some neurological disorders such as AD, traumatic brain injury, and vascular dementia. Moreover, this spatial testing paradigm can be utilized to detect more subtle cognitive impairments in experimental animals which may not show impairments during original location training.

### Novel findings in the current report

The development of the 3-phase variant of the MWT for mice resulted in several new findings. First, a statistically significant amount of extra training was required to get the quantitative and qualitative levels of spatial learning and memory exhibited by rats using standard training procedures. Second, the use of various measures of place learning and memory provided clear evidence of precise representations acquired by mice when more extensive training procedures were instituted. Finally, we showed that mice could acquire multiple representations about the location of escape platforms in the same apparatus and context. A brief exploration of each of these novel findings and their potential implications will be discussed below.

The protocol for MWT varies from laboratory to laboratory depending on the desired end points of the particular study. Despite the extensive application of the MWT in behavioural neuroscience to test spatial learning and memory in rodents, the optimal methodological parameters for the use of this paradigm in mice is not clear. Different research groups follow different protocols for MWT for mice without stating the importance of the change in parameters^34-36^. For example, some research groups used MWT training parameters in mice with either 2-4 trials per day for 4-5 days along with a probe trial on either fifth or sixth day^34-37^. In our hands, this would not have been a sufficient amount of training to reach asymptotic levels of performance and precise spatial navigational abilities. One reason why this can be problematic is because this suboptimal level of performance on the task can lead to erroneous conclusions about the neural circuits involved^38^ and the neurobiological mechanisms required^19^.

The addition of rapid mass training in on a session to a new platform location in this mouse paradigm is unique. In previous studies, researchers assessed new location learning in mice for 5-6 days for which is similar to the normal acquisition procedures except the location of the platform^27,33^. Furthermore, in these studies, they do not perform a competitive probe trial to distinguish the influence of the old and new memory representations on the behaviour of the mice. To overcome these drawbacks and in order to optimize the procedure for assessing the old and new spatial memory of mice within MWT, we included a rapid mass training phase to assess how quickly the mice will learn the new location of the platform and when the new representation would gain control over voluntary behavior. We showed that with the right training parameters, mice could exhibit knowledge of two spatial representations of the platform location.

One surprising and fascinating finding from the current set of experiments was that *App*^*NL G-F/NL-G-F -*^ mice showed severe learning and memory impairments when trained over multiple days but during a rapid mass training day showed the ability to learn a new spatial location. This pattern of results suggests that the impairment exhibited by these mice is linked to alterations encoding mechanisms that can be overcome with intensive training or in memory decay or retrieval mechanisms. Consistent with the latter idea, human research and mouse models of AD suggest the memory impairment in early AD might be due to a failure to retrieve encoded information rather than an inability to encode that information^39^.

The MWT is an elegant and versatile task for assessing spatial learning and memory functions of rodents. Many procedural variations of the MWT have been and are being used by research groups in many different applications. The new protocol employed in the present experiments provides an important addition for use in the mouse with various advantages over other designs.

### Other examples of the potential utility of the 3-phase variant of the MWT for research using rat and mouse models?

The 3-phase variant of the MWT explored in the current experiments has statistically significant advantages over the standard version of the task for investigations into the neural basis of learning and memory in the rodent, studies of the neurobiological mechanisms supporting learning and memory functions, and the assessment of use of animal models of human disease states like AD.

One key advantage of this version of the MWT is that it allows the ability to compare the effects of experimental brain manipulations during acquisition of spatial information in a naive versus procedurally sophisticated subjects. This is important because, for example, several key mechanisms of plasticity processes thought to support hippocampal learning and memory functions have been questioned because the effects occur in naive subjects but not procedurally sophisticated subjects^19^. How does the processing style of the hippocampus relate to the importance of the 3-phase variant of the MWT? We have argued that this distributed nature of hippocampal memory representations makes the learning and memory system centered on the hippocampus difficult to study when the experimental manipulation employed by the researcher has less than complete effects on the entire vertical and horizontal expanse of the structure. This is a particular problem when the researcher only manipulates a part of the hippocampus and reports particular effects of this manipulation as characteristic of the key functions of the system. These studies suggest that animals can display intact memory in these tasks despite substantial but incomplete damage to the HPC. In fact, we have argued that the standard version of the MWT might be insensitive to this type of incomplete hippocampal dysfunction and there is support for this claim in the literature.

The 3-phase variant of the MWT developed in the current report, therefore, is important for the kind of work described above in which partial hippocampal dysfunction is possible and the need for a more sensitive assay of hippocampal function. We believe that the 3-phase version of the MWT is also important because the new location training component of the paradigm is thought to place a higher demand on hippocampal processing^40^.

### Summary

A variant of the MWT for use in mice was explored. The task consists of three phases including original location training, rapid new location training, and a competition test between the old and new location quadrants. Although originally developed for use in rats, we found that mice can show similar learning abilities on the different phases of the task including the ability to acquire and store multiple location memories with some procedural changes. It was clear that more extensive training was required to get the kind of precise place learning and memory abilities of the rat and it is important to use a variety of dependent measures of this complex spatial navigation ability. The results showed that C57 mice acquired and retained both the old and new location quadrants associated memory, conversely, *App*^*NL-G-F/NL-G-F*^ mice reserved only a recently acquired spatial memory but did not retain the memory with the old platform location acquired in the same apparatus and context. The results also indicated that C57 mice can show precise place learning and memory with the right amount of training and acquire and retain multiple spatial memory locations in the same environment whereas this ability was compromised in *App*^*NL-G-F/NL-G-F*^ mice. It is argued that this task can be used in various basic and applied approaches to understand the neural basis of learning and memory, the key role of the hippocampus in these processes, the etiology of age-related dementia, and development of preventative and treatment approaches for these devastating neurodegenerative diseases.

## Acknowledgements

This work was supported by Natural Sciences and Engineering Research Council of Canada (NSERC) Discovery Grant #40352 and #06347 to MHM and RJM respectively, Campus Alberta for Innovation Program Chair (MHM), Alberta Alzheimer Research Program (MHM & RJM), Alzheimer Society of Canada (MHM & RJM). We thank Di Shao and Behroo Mirza Agha for animal breeding.

